# The lipidome and proteome of high-density lipoprotein are altered in menopause

**DOI:** 10.1101/2024.01.10.574516

**Authors:** Satu Lehti, Tia-Marje Korhonen, Rabah Soliymani, Hanna Ruhanen, Emilia Lähteenmäki, Mari Palviainen, Pia Siljander, Maciej Lalowski, Reijo Käkelä, Maarit Lehti, Eija K Laakkonen

## Abstract

High-density lipoprotein particles (HDL) possess anti-inflammatory, anti-thrombotic, cytoprotective, and anti-oxidative properties, thus protecting against cardiovascular diseases. Menopause is associated with changes in serum metabolome and HDL size distribution. We analyzed the protein and lipid composition of the HDL particles from pre-, peri-, and postmenopausal women (N=216) with nuclear magnetic resonance and mass spectrometry to get a deeper insight into the composition of HDL in different stages of menopause. Both particle size and composition differed; in perimenopause, the proportion of small HDL particles (8.7 nm on average) was higher, and the proportion of large HDL particles (12.1 nm on average) was lower than in pre- or postmenopause. In perimenopause, each particle size class was enriched with triacylglycerols, and the calculated lipid class ratio of triacylglycerol/cholesteryl ester was the highest within perimenopausal HDL particles. This potentially affects the HDL interaction with lipid-modifying enzymes. We also observed directionally opposite associations for HDL cholesteryl ester and unesterified cholesterol with systemic estradiol and follicle-stimulating hormone levels, especially regarding S-sized HDL particles, but not the hormone associations with HDL triacylglycerols. Perimenopausal HDL also exhibited a lower proportion of apolipoproteins (apoA-I, apoA-II, apoC-I, apoC-III, apoD and apoE) per particle than premenopausal or postmenopausal HDL. In summary, we found that premenopausal and postmenopausal HDL particles were compositionally similar and differed from perimenopausal ones. We suggest that menopause, and especially the unbalanced hormonal state in perimenopause, are reflected in the lipid and protein compositions of the HDL, which, in turn, may affect the functions of the HDL particle.

## Introduction

Menopause affects female aging in multiple ways as reflected by the circulating metabolome. Our research, along with others, has demonstrated significant changes in serum lipid metabolome towards the proatherogenic direction (Auro et al., 2014; Karppinen et al., 2022; Wang et al., 2018). The high-density lipoprotein cholesterol (HDL-C) tends to rise from premenopause to postmenopause (El Khoudary et al., 2021) and HDL particle size distribution shifts towards smaller diameters (Karppinen et al., 2022; Wang et al., 2018). Furthermore, a net cholesterol efflux capacity per HDL particle temporally decreases during menopausal transition (El Khoudary et al., 2021). That prompted the authors to suggest that HDL functioning may deteriorate at menopause, which would explain why increasing HDL-C concentrations occur concurrently with worsening cardiovascular health during the menopausal transition.

In circulation, HDL particles form a heterogeneous population, varying from small discoidal ones to larger spherical particles (sizes 7-12 nm). In addition to a variety of lipids (Kontush et al., 2013), HDL particles contain a versatile mixture of proteins (Davidson et al., 2022), of which apolipoprotein (apo) A-I and apoA-II are integral structural proteins. This compositional heterogeneity allows HDL particles to display a great functional variety. HDL particles have, for instance, been found to modulate functions of innate immunity, glucose metabolism, and cellular communication (Von Eckardstein et al., 2023). However, a detailed characterization of HDL lipidome and proteome across different menopause statuses is currently lacking.

Our study was motivated by the desire to determine whether structural differences exist between pre-, peri- and postmenopausal HDL. We investigated the HDL lipidome and proteome using nuclear magnetic resonance (NMR), mass spectrometry (MS), and gas chromatography (GC) to characterize the molecular structure of the particles derived from pre-, peri-, and postmenopausal women. NMR enabled size-based categorization of serum HDL particles and revealed the distribution of major lipid classes within each HDL particle size group. Subjecting the isolated HDL particles to MS enabled both the characterization of the lipid species and the proteome of the particles while GC provided further insights into the HDL lipidome by allowing fatty acid level analysis. We hypothesize that the hormonal turmoil during menopause is reflected in the composition of HDL, which may, in turn, affect HDL functionality.

## Results

To assess the compositional differences and functional properties of HDL particles between menopausal statuses, we selected 216 healthy, non-medicated normolipidemic women aged 47–56 years from the Estrogenic Regulation of Muscle Apoptosis (ERMA) cohort (Kovanen et al., 2018) representing premenopause (PRE), perimenopause (PERI) and postmenopause (POST) statuses. The selected women were not using hormonal contraception, menopausal hormone therapy, or medication affecting lipid metabolism. The menopausal status of each woman was determined by hormone measurements and the menstrual diaries provided by the women. To minimize within-group variation among individuals and to obtain sufficient serum sample volume, allowing several analyses from the same isolated HDL fraction, six individual serum samples were combined into a pool, and 12 such pools per menopausal stage were used in the analyses (Figure 1). Characteristics of the three menopause groups are presented at the group means level in Table 1 and as a pooled sample level in Table S1. Menopausal groups were otherwise similar except that women in the POST group were about two years older and had lower estradiol and sex hormone-binding globulin and higher follicle-stimulating hormone (FSH) levels demonstrating that the menopause had occurred. In addition, the clinical measures of total cholesterol, low-density lipoprotein cholesterol (LDL-C), and HDL-C were the highest, and glucose was the lowest in the POST group. Despite small group differences, the body mass index (BMI) of each pool was close to 25, with very similar ranges across menopausal groups (PRE 24.7–26.0, PERI 24.6–26.1, POST 24.2–26.0).

**Table 1.**
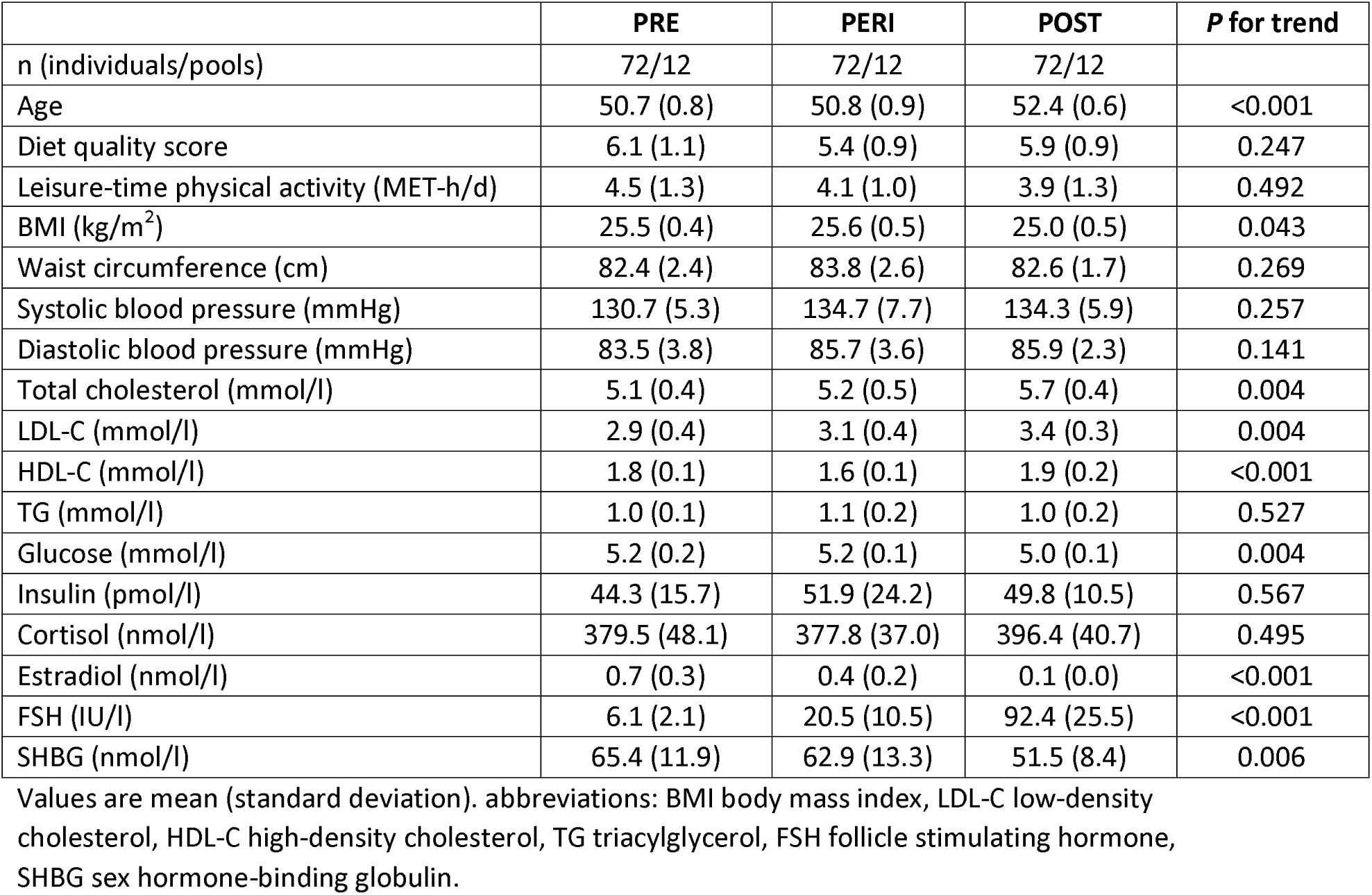
Characteristics of the menopausal groups.

**Figure 1.**
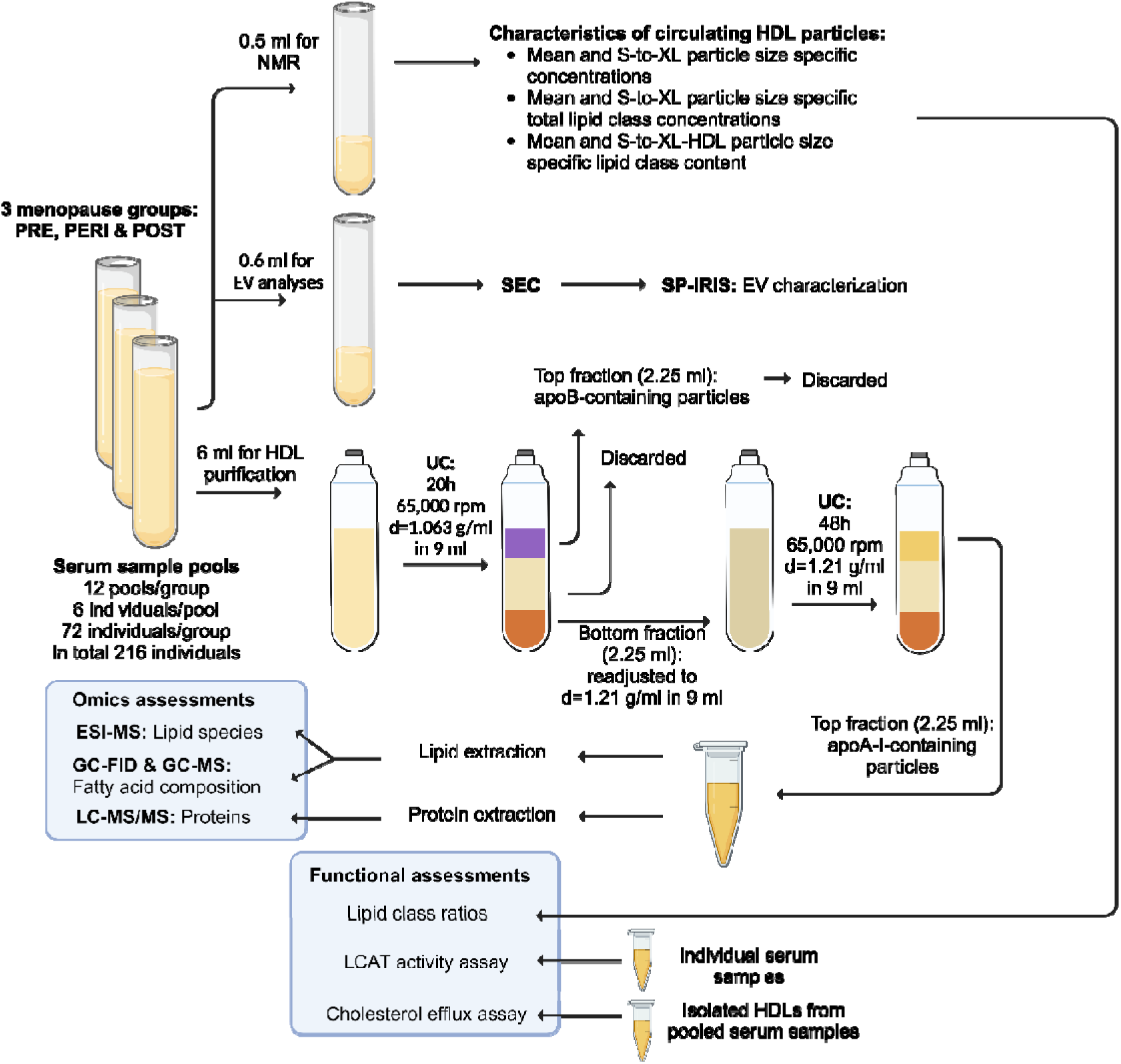
Schematic of the study design and sample workflow. Abbreviations: PRE, premenopausal; PERI, perimenopausal; POST, postmenopausal; NMR, nuclear magnetic resonance; HDL, high-density lipoprotein particle; EV, extracellular vesicle; SEC, size exclusion chromatography; SP-IRIS, single particle interferometric reflectance imaging sensing; UC, ultracentrifuge; apoB, apolipoprotein B; apoA-I, apolipoprotein A1; ESI-MS, electrospray ionization-mass spectrometry; GC-FID, gas chromatography with flame ionization detector; GC-MS, gas chromatography-mass spectrometry; LC-MS/MS, liquid chromatography-tandem mass spectrometry; LCAT, lecithin cholesterol acyltransferase.

### Particle characteristics of pre-, peri- and postmenopausal HDL

The average HDL particle sizes and concentrations, the concentrations of small (S), medium (M), large (L), and extra-large (XL) -sized HDL particles (8,7 nm, 10.9 nm, 12.1 nm, and 14.3 nm in average, respectively), as well as the total lipid concentration, and main lipid class concentrations of the particles were profiled from pooled serum samples using an NMR analysis platform focused on lipoproteins and lipids (Soininen et al., 2015). The mean HDL particle size (Figure 2a, PRE - PERI p = 0.014, PERI - POST p = 0.029) and its concentration in serum (Figure 2b, PERI - POST p = 0.0006) were measured to be the lowest in the PERI group. The S-HDL particles constituted 62.3% (CI 61.1-63.4) of all particles in PERI, and 59.6% (CI 58.4-60.7) and 59.9% (CI 58.2-61.5) in PRE and POST groups, respectively (Figure 2c). Consequently, in the PERI group the S-HDL lipid content was higher (Figure 2d, PRE - PERI p = 0.015, PERI - POST p = 0.030), while in the L-HDL the lipid content of PERI was lower (PRE - PERI p = 0.024, PERI - POST p = 0.038) than in PRE or POST groups. There were no differences between menopausal groups regarding M- or XL-particles.

**Figure 2.**
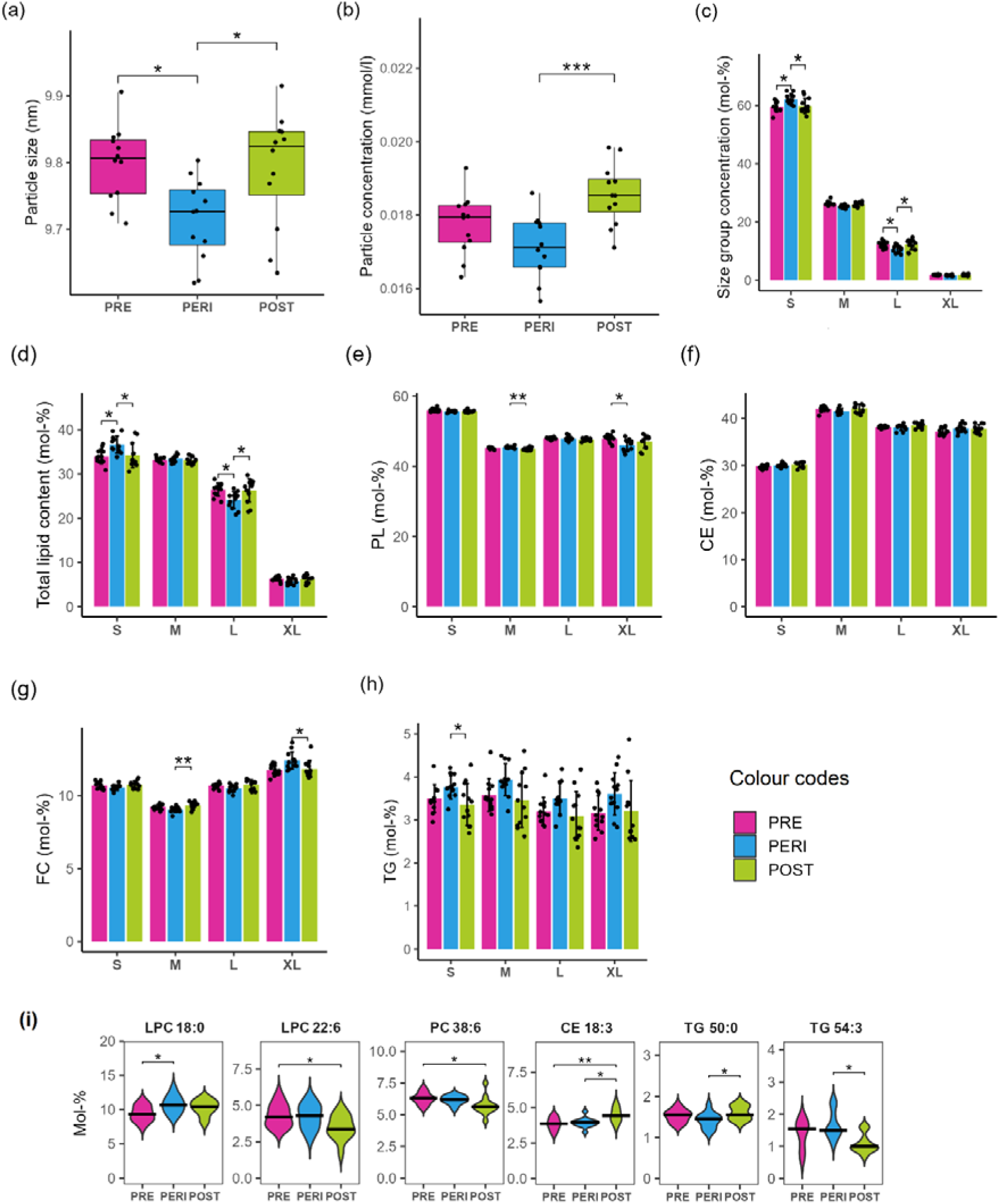
The main lipid classes and the species of HDL particles in pre-, peri- and postmenopausal women. (a) Average HDL particle sizes in menopause groups. (b) Average particle concentration in serum of menopausal women. (c) HDL particle concentrations according to their size categories within each menopausal group. (d) Total lipid content of the HDL particles in each size category. (e) Size category-specific relative concentration of phospholipids. (f) Size category-specific relative concentration of cholesteryl esters. (g) Size category-specific relative concentration of unesterified cholesterols. (h) Size category-specific relative concentration of triacylglycerols. (i) The lipid species of the HDL particles, which possess statistically significant differences in their relative concentration between menopausal groups. Statistical significance *<0.05, **<0.01, ***<0.001. Abbreviations: PRE, premenopause; PERI, perimenopause; POST, postmenopause; HDL, high-density lipoprotein; PL, phospholipid; CE, cholesteryl ester; FC, unesterified cholesterol; TG, triacylglycerol; LPC, lysophosphatidylcholine; PC, phosphatidylcholine.

The molar percentages of the four major lipid classes: phospholipids (PL), cholesteryl esters (CE), unesterified cholesterols (FC), and triacylglycerols (TG) are presented for each HDL particle size group in Figure 2e-h. Subtle group differences were observed in the proportions of PLs (Figure 2e, M-HDL was higher in PERI than POST group, p = 0.006 while XL-HDL was lower in PERI than PRE group, p = 0.026) and FCs (Figure 2g, M-HDL was lower in PERI than POST group, p = 0.003 and XL-HDL was higher in PERI than POST group, p = 0.025). The PERI group displayed the highest proportion of TGs, especially in the S-HDL particles (Figure 2h, higher in PERI than POST group, p = 0.033).

### Characterization of serum extracellular vesicles as a potential source of bias for the ultracentrifugally isolated HDL

For in-depth HDL lipidomic and proteomic analyses, the HDL particles were isolated using sequential ultracentrifugation (Figure 1). However, it is known that thusly separated HDL (d<1.21 g/ml) may also contain extracellular vesicles (EVs) (Simonsen, 2017). To evaluate if menopausal groups differed in their EV content, which could then constitute a significant source of bias towards our HDL-omics results, we utilized size exclusion chromatography (SEC) to isolate EVs from aliquots of the same pooled serum samples subsequently used for HDL-omics analyses. The isolated EVs were characterized using ExoView technology with anti-CD41, -CD9, -CD63, and -CD81 immunostaining. The sizes of EVs were similar in all menopausal groups, and we observed no statistical differences in the PRE, PERI, and POST groups regarding the distribution of tetraspanins CD81 and CD63 (Figure S1). However, there was a statistically significant difference between the PRE and POST groups regarding EV markers relevant for platelet-derived EVs (CD41 and CD9, p = 0.002 and 0.013, respectively), suggesting that the POST serum samples exhibited reduced levels of platelet EVs compared to PRE group. Neither group differed from the PERI group. We considered the identifications of EV-derived lipids and proteins to be negligible compared to the more numerous HDL-derived hits (Nieuwland & Siljander, 2023).

### HDL lipidomics: lipid species and fatty acids

Following HDL isolation by sequential ultracentrifugation, the HDL lipids were extracted, and lipid species were analyzed with electrospray ionization-mass spectrometry (ESI-MS) (Figure 1). From the total lipid extract, fatty acids (FA) were derivatized into fatty acid methyl esters (FAMEs) and analyzed with gas chromatography using flame ionization detectors (GC-FID) and with combined gas and mass chromatography using mass-selective detectors (GC-MS).

The lipid species with a relative concentration over 0.5 mol-% were included in the main analysis. For each lipid class, we present the six most abundant lipid species in Supplementary Figure S2. In all menopausal statuses, HDL lipid species composition followed the same pattern as observed in previous research (Kontush et al., 2013). Lysophosphatidylcholine (LPC) 18:2 and LPC 16:0 together accounted for over 50 mol-% of all LPCs. The most abundant phosphatidylcholine (PC) was PC 34:2 accounting for about 24 mol-% of all PCs. CE 18:2 was the most abundant CE species (64 mol-% of all CEs). Together sphingomyelin (SM) 34:1;O2 and SM 42:2;O2 accounted for 47 mol-% of the SMs. In this dataset, the most abundant TG was TG 50:1 with about 20 mol-% portions of all TGs. We also observed some significant group differences (Figure 2i) for LPC 18:0 (highest in PERI, lowest in PRE; PRE - PERI p = 0.025), LPC 22:6 (highest in PRE, lowest in POST; PRE - POST p = 0.049), PC 38:6 (highest in PRE, lowest in POST; PRE - POST p = 0.040), CE 18:3 (highest in POST, lowest in PRE; PRE - POST p = 0.004, PRE - PERI p = 0.014), TG 50:0 (highest in POST, lowest in PERI; PERI - POST p = 0.034) and TG 54:3 (highest in PERI; lowest in POST; PERI - POST p = 0.015).

For FA analysis, species with relative concentrations above 0.1 mol-% were included. The highest mol-% of odd-chain FAs 15:0, 17:0 *anteiso* (a branched chain FA), and 17:0 (p = 0.008, p < 0.001 and p = 0.018, respectively) were observed in the PRE, medium values in the PERI and the lowest relative concentrations in the POST group. Significant group differences were detected between the PRE and POST statuses (Figure S3a). For the polyunsaturated FAs of the n-6 pathway, the lowest relative concentrations were found in the PRE and the highest in the POST group. Of them, only the intermediary FA 18:3n-6 differed significantly between the groups (p= 0.022) (Figure S3b).

To summarize the lipid results, although there were differences in the lipid classes between the menopausal statuses, only six lipid species differed statistically significantly. Most of the FAs found to differ were odd-chain FAs, and only one even-chain FA was found to differ between the menopausal statuses.

### HDL proteomics

Proteomic analysis utilizing liquid chromatography-tandem mass spectrometry (LC-MS/MS) was carried out on the same ultracentrifugally isolated HDL particles as the lipidomics analysis (Figure 1). Of the total 149 proteins that were detected in menopausal HLD, 44 proteins were preliminarily quantified based on being present in at least two out of three repeated injections of the same sample. The proteins were annotated by their known functions within Gene Ontology (GO) molecular function or biological processes terms mapping into four major functional classes including lipid metabolism, transport and signaling, immune defense, and regulation of cellular processes (Figure 3a and Table S2). By applying more stringent filtering criteria, 19 proteins remained to be simultaneously detected in eight out of twelve HDL samples per each menopausal group.

**Figure 3.**
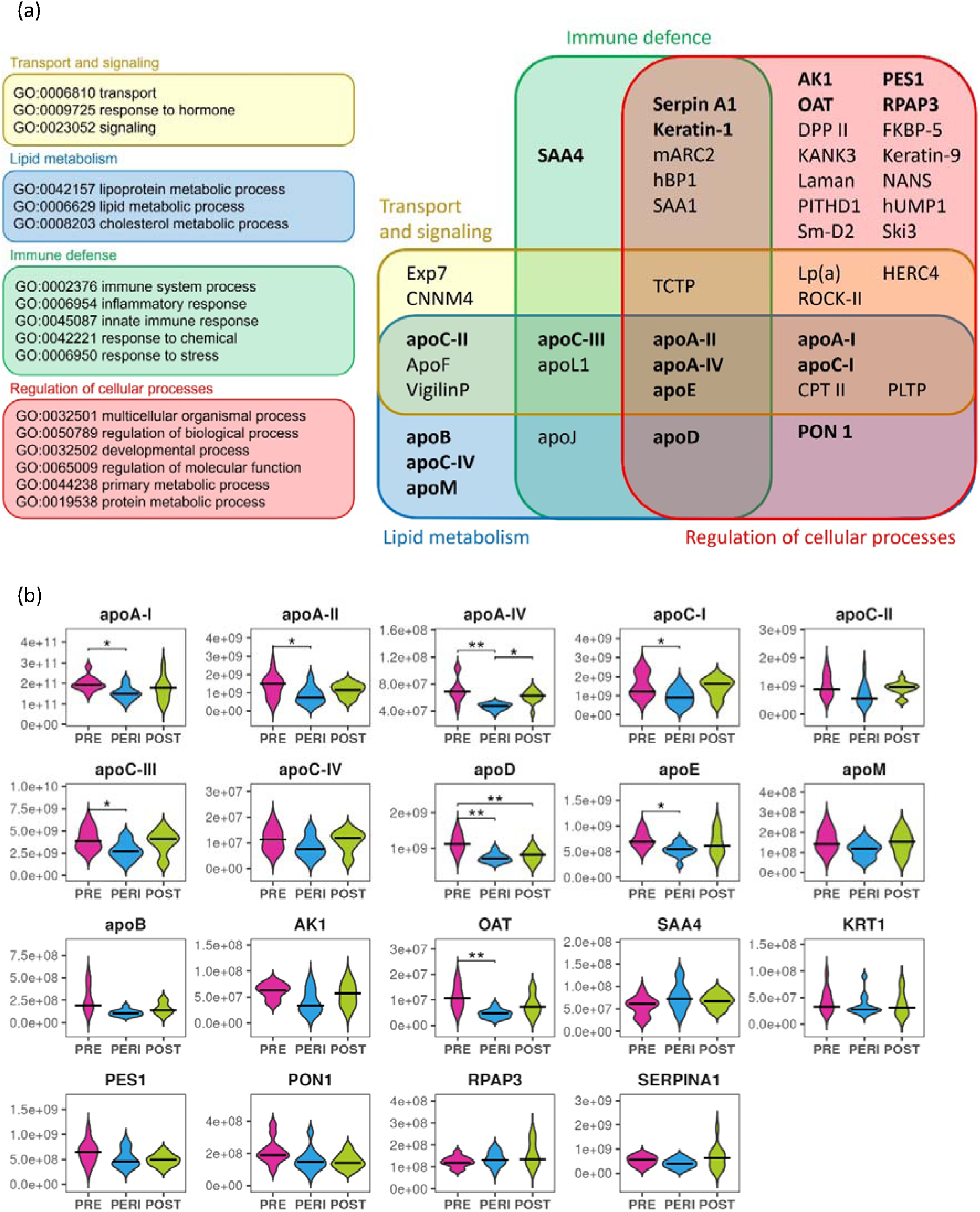
Proteomic classification of HDL particles in menopause. (a) The 44 quantified HDL proteins were associated with four main categories of Gene Ontology. The 19 proteins, which were simultaneously detected in eight out of twelve HDL samples per menopause group are bolded. (b) Violin plots of HDL protein abundances normalized to NMR-derived HDL particle count. All protein abundances were measured with the LC-MS/MS method from the isolated HDL particles. Statistical significance * < 0.05, ** < 0.01. Abbreviations: PRE, premenopause; PERI, perimenopause; POST, postmenopause; HDL, high-density lipoprotein; LC-MS/MS, liquid chromatography-tandem mass spectrometry; NMR, nuclear magnetic resonance.

Since we observed differences in HDL particle counts between the menopausal groups (Figure 2b), we present here the HDL protein abundances normalized to NMR-derived HDL particle counts (Figure 3b). The particle counts were measured from the same pooled serum samples from which the HDL particles were isolated. The non-normalized data are presented in Figure S4. HDL particles may contain multiple copies of apoA-I, with up to five copies in the largest particles (Malajczuk et al., 2021). Therefore, we also calculated the protein abundances normalized to apoA-I, measured simultaneously with LC-MS/MS (Figure S5). Interestingly, 14 out of the 19 quantified proteins (normalized to HDL particle count) exhibited the lowest relative amount in the PERI compared to PRE or POST stages, although not all differences were statistically significant (Figure 3b).

Apolipoproteins are the most abundant protein group of HDL. Among them, apoA-I and apoA-II are the main structural apolipoproteins of HDL. All apolipoproteins had their lowest abundances in the PERI samples even when the lower number of HDL particles in the PERI group was considered (Figure 3b). Ratio of apoA-II to apoA-I and the ratio apoA-II to HDL particle counts were the lowest in the PERI samples, suggesting that either there were fewer apoA-II apolipoproteins in each HDL particle or fewer apoA-II-containing HDL particles.

Of the proteins classified with immune defense and/or cellular regulatory functions, the PERI group displayed, as compared to the PRE, a significantly lower relative abundance of adenylate kinase isoenzyme 1 (AK1) and ornithine aminotransferase (OAT) (Figure 3b). In contrast, the relative abundance of serum amyloid A-IV (SAA4), which is a constitutively translated apolipoprotein linked to inflammatory diseases, was higher in the PERI compared to the PRE, but the result reached statistical significance only when normalized to apoA-I abundance (p = 0.015) (Figure S5).

To conclude, most of the proteins, and notably, all apolipoproteins, exhibited their lowest abundance (non-normalized or normalized to HDL-P) in PERI group and concomitantly, the highest abundance in PRE group. The few exceptions for these trends were serum paraoxonase (PON1) and alpha-1-antitrypsin (Serpin A1) with decreasing trends across menopausal statuses from pre-to-postmenopause, and RNA polymerase II-associated protein 3 (RPAR3) and SAA4 with increasing trends from pre-to-postmenopause.

### Multi-omics integration analyses

Next, we used three-dimensional partial least squares discriminant analysis (3D-PLS-DA) to inspect if combining omics results grouped into one class instead of a single omics item would separate menopausal statuses from each other. The analytical entities for the 3D-PLS-DA were combined PC and LPC classes, SMs and TGs, main lipid classes together, FAs and proteins (Figure 4a). The clearest separation of the three menopausal groups was observed with the main lipid classes and proteins.

**Figure 4.**
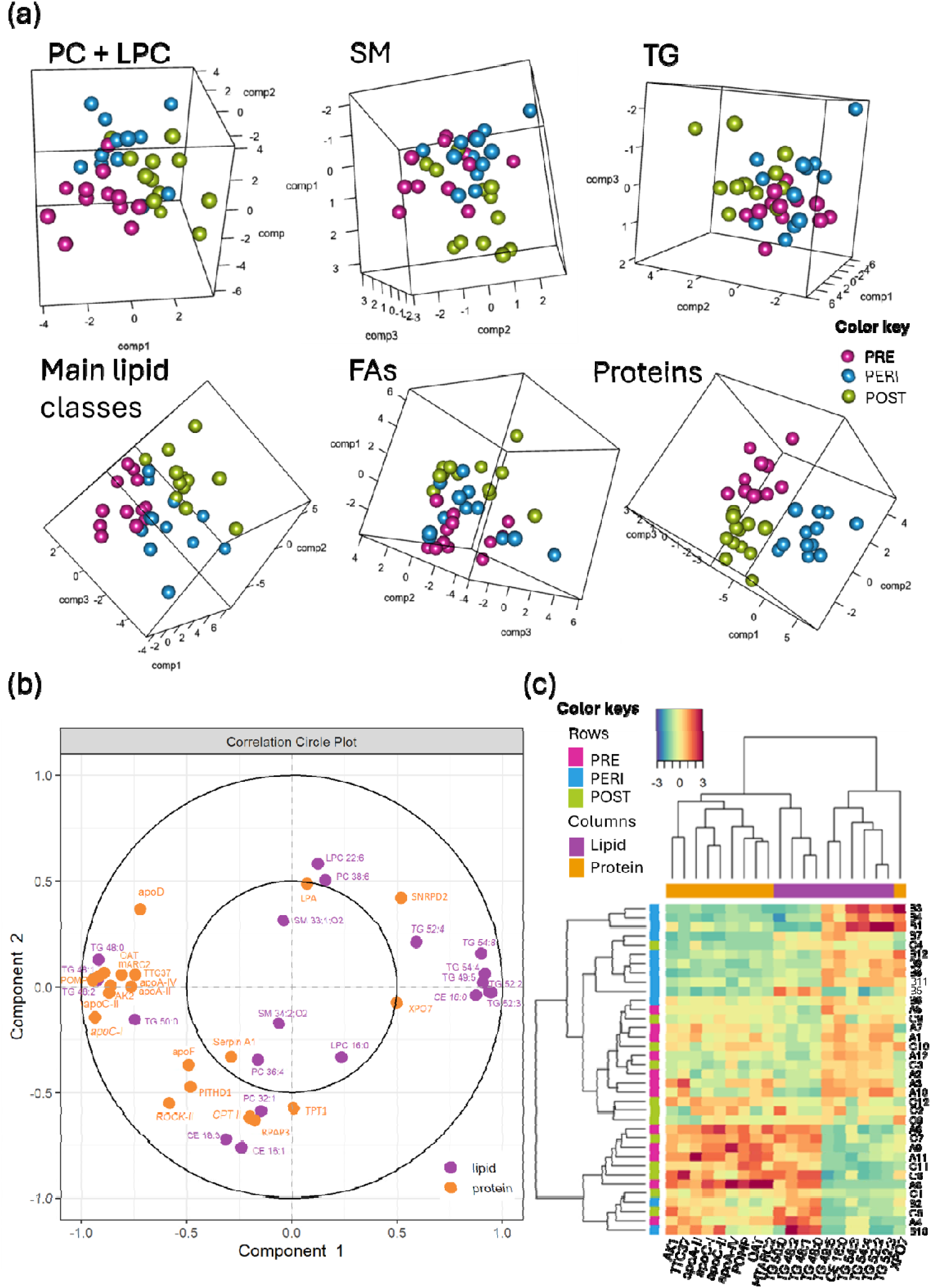
Multi-omics analyses of HDL lipids and proteins. (a) PLS-DA scores plots for combined phosphatidylcholine (PC) and lysophosphatidylcholine (LPC), sphingomyelin (SM), triacylglycerol (TG), all main lipid classes, all fatty acids (FAs) and all proteins. Lipid species and fatty acids were analyzed by non-targeted mass spectrometry and gas chromatography, respectively. Quantified protein profiles were assessed with non-targeted liquid chromatography-tandem mass spectrometry. (b) Unsupervised correlation circle plot of analyzed lipid species and proteins. No correlation cut-off was applied. (c) Heat map of lipid and protein abundances. Y-axis represents individual serum pools. As inputs to PLS-DA, 44 proteins and 87 lipid species were used, and those that most significantly differentiated menopausal statuses were depicted in the correlation circle (b) and the heat map (c). Abbreviations: PRE, premenopause; PERI, perimenopause; POST, postmenopause; PC, phosphatidylcholine; LPC, lysophosphatidylcholine; SM, sphingomyelin; CE, cholesteryl ester; TG, triacylglycerol; FAs, fatty acids.

The correlation plot shows the relationship between variables, in this case lipids and proteins of HDL particles. Here, all 44 proteins were loaded as variables. In the correlation circle, objects on the opposite sides of the circle indicate a negative correlation between these two objects, while objects close to each other represent the positive correlation. Figure 4b shows that apoA-I, apoA-II, apoA-IV, apoC-I, and apoC-II, as well as mitochondrial proteins OAT and mARC2 positively correlate with saturated and mono- and diunsaturated TGs (48:0, 48:1, 48:2, and 50:0). They correlated negatively with polyunsaturated TGs (49:4, 52:2, 52:3, 52:4, 54:4, and 54:8). The unsaturated TGs correlated also positively with CE 16:0 which was found in the same tightly packed cluster. CE 18:3 and CE 18:1 showed no correlation with the above-mentioned lipids or proteins, while they positively correlated with each other, PC 36:4 and 32:1, and Serpin A1. They also correlated negatively with PC 38:6, LPC 22:6, and SM 33:1;O2. In the heatmap presented in Figure 4c, the PERI samples clustered together, while the PRE and POST samples appeared intermingled (rows), and lipids and proteins formed separate clusters (columns). Interestingly, many of the left-side cluster proteins of the circle plot were also included among the protein cluster of the heatmap, thereby strongly contributing to the separation of the PERI samples from the other menopausal groups.

### Functional characterization of HDL within menopausal groups

We took several approaches to gain insight into whether menopause-associated differences in the HDL composition could manifest as functional differences. First, we calculated ratios of lipid classes, which can be used as indicators of the function of HDL-linked enzymes such as cholesteryl ester transfer protein (CETP) and lecithin cholesterol acyltransferase (LCAT). Ratios were calculated using the NMR data of major lipid classes and HDL particles and are presented as HDL size group-specific values (Figure S6). The higher TG/CE ratio reflects higher CETP activity as CETP transfers and exchanges CE and TG between lipoproteins. TG/CE ratio (Supplementary Figure S6a) was the highest in PERI in every HDL size group, although significant only in XL-HDL (PERI - POST p = 0.048). The activity of the LCAT, which esterifies FC on the surface of HDL to form CE (the esterified FA is derived from PC), can be estimated using ratios FC/PL and FC/CE (higher LCAT activity leads to lower ratios) The statistically significant group differences for FC/PL (Figure S6b) were observed in M-HDL (PERI - POST p = 0.002), where PERI group displayed the lowest ratio, and in XL-HDL (PRE - PERI p = 0.013), in which the PERI group exhibited the highest ratio. The ratio FC/CE (Figure S6c) differed significantly only for XL-HDL, where PERI group showed a higher value than PRE (p = 0.040) and POST (p = 0.007).

Second, we assessed if the lipid ratios in the pooled serum samples could be linked to actual differences in serum LCAT activity in individual samples representing the different menopausal groups. Therefore, we measured the LCAT activity of ten PRE, nine PERI, and nine POST women. To avoid whole-body adiposity-related differences, each participant’s BMI was matched with each other in addition to the pooled BMI of the menopausal groups. The mean BMI was 25 ± 0.3 kg/m^2^. We did not observe differences in the measured LCAT activity between the groups (Figure S6d). LCAT activity, however, was found to correlate negatively (r = -0.475, p = 0.011) with serum estradiol concentration. Interestingly, the correlation was not linear at the low estradiol range; the women with low estradiol concentration, mainly representing the POST group, had a high variation with LCAT activity, but not in estradiol concentration (Figure S6e).

Third, to find out if the menopausal status is associated with differences in the cholesterol efflux capacity of the HDL, we measured the ability of the isolated HDL from the different menopausal groups to take up fluorescently labeled cholesterol from THP-1-monocyte-derived macrophages. Although a slight increase in the efflux capacity was observed across advancing menopausal status, no statistical significance between the groups was detected (Supplementary Figure S6f).

### Association of estradiol and FSH with HDL particle characteristics

To assess the potential role of menopausal hormonal differences, we inspected the associations of estradiol and FSH on measured HDL particle characteristics using linear regression models controlled for confounders of waist circumference and leisure-time physical activity (all models are presented in Table S3). We found negative associations with estradiol concentration and positive associations with FSH concentrations for HDL-PL, -CE, and -FC (Figure 5a). Regarding estradiol, the associations were stronger with smaller HDL particles reaching statistical significance for S-HDL-CE (β = -0.47 ± 0.14, p = 0.003) and S-HDL-FC (β = -0.49 ± 0.15, p = 0.003). FSH, of which level rises more steadily than estradiol falls during approaching menopause, seemed to be a stronger predictor of HDL-lipids than estradiol. The highest standardized regression coefficient estimates were observed for L-, M- and S-sized HDL particles with stronger statistical evidence for S-HDL-CE (β = 0.55 ± 0.14, p < 0.001) and S-HDL-FC (β = 0.58 ± 0.15, p < 0.001). HDL-TG was not associated with estradiol or FSH.

**Figure 5.**
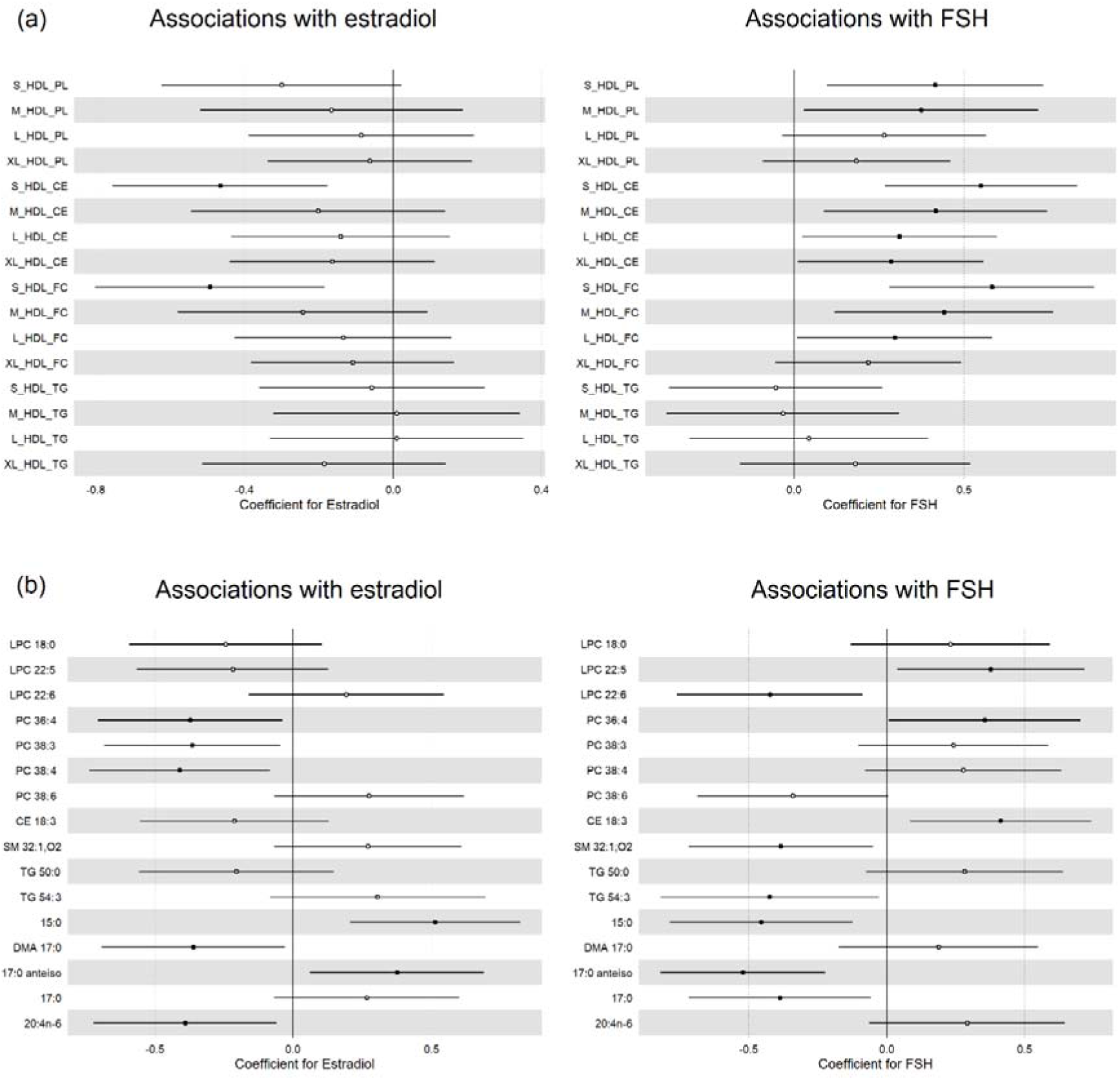
Associations of estradiol and FSH with HDL particle characteristics. (a) Associations of estradiol and FSH with HDL particle size category-specific relative concentrations of main lipid classes measured by nuclear magnetic resonance spectrometry. (b) Associations of estradiol and FSH with lipid species and fatty acids of the HDL particles measured by mass-spectrometry and gas chromatography. The linear regression models were controlled with waist circumference and leisure time physical activity. Significance p<0.05 marked with closed circles. Abbreviations: FSH, follicle-stimulating hormone; HDL, high-density lipoprotein; PL, phospholipid; CE, cholesteryl ester; FC, unesterified cholesterol; TG, triacylglycerol, LPC, lysophosphatidylcholine; PC, phosphatidylcholine; SM, sphingomyelin.

Similarly, opposing trends were observed for the associations of estradiol and FHS with HDL-carried lipid species and their fatty acid composition (Figure 5b, Table S4). Among the lipid species differentially expressed between menopausal groups (Figure 2), only LPC 22:6 (β = -0.42 ± 0.17, p = 0.015) and CE 18:3 (β = 0.41 ± 0.16, p = 0.015) were significantly associated with FSH, but none with estradiol. We also constructed regression models for the other lipid species as well as for the fatty acids presented in Figure S3 and plotted them in Figure 5b when notable associations were found for estradiol or FSH. Four odd-chain FAs (15:0, 17:0, plasmalogen dimethyl acetal (DMA) 17:0, and a branched chain FA 17:0 *anteiso*), as well as arachidonic acid 20:4n-6 showed significant associations with estradiol albeit they were not statistically significantly different between the menopausal statuses. Also, a few PC-, LPC-, and SM-species were found to be associated with estradiol (PC 36:4, 38:3, and 38:4) or FSH (LPC 22:5, PC 36:4, and SM 32:1;O2), although the species were not found to differ significantly between the menopausal statuses.

## Discussion

In this study, we performed an in-depth analysis of HDL from women representing three menopausal statuses to investigate menopause-associated differences in the HDL particle characteristics (size, lipidome, and proteome). In our cross-sectional study, we observed rather unexpectedly that, in many of the studied variables, the perimenopausal HDL profile stood out from premenopausal and postmenopausal profiles. It appeared that in perimenopause, the average particle size and concentration were lower than in pre- or postmenopausal women. The concentration of small HDL particles, the total lipid and TG content of particles, as well as the ratio of TG to CE, were higher in perimenopausal women’s HDL compared to pre- or postmenopausal women’s HDL. The proteomic analysis showed that many apolipoproteins had lower abundances in the perimenopausal women’s HDL than in pre- or postmenopausal women. The FA analysis broke the pattern; FA results followed the expected sequence, showing systemic increment or decrement from pre- to postmenopause, while significant differences in HDL-associated lipid species with the highest or lowest abundances were observed for all menopausal groups. The constructed regression models further highlighted the importance of menopausal hormones for HDL lipid content. For instance, we observed that lower estradiol and higher FSH levels were associated with higher HDL-CE and HDL-FC, especially in S-sized particles, but not with HDL-TG.

The HDL lipid composition largely dictates the particle size, which in turn influences protein composition by determining how versatile protein contents fit within the particle. That may ultimately determine the functional properties of HDL in different menopausal statuses. In our study, TG proportions were the highest in perimenopausal HDL, as were the proportions of PL in the M-sized, and lowest in the XL-sized HDL particles, whereas Wang *et al*. (Wang et al., 2018) measured low HDL-TG and -PL in postmenopausal women’s sera. Others have also concluded that diminished estrogen availability in peri- and postmenopause may decrease HDL particle size and change its PL composition (Piperi et al., 2004; Tangney et al., 2001). This is important, because small size, low PL, and high TG contents of HDL are associated with cardiovascular diseases (Kontush et al., 2013).

Further insight into HDL’s functional properties can be gained by inspecting the ratios of lipid classes, which can indirectly reveal the efficiency of different lipid-associated enzymes. CETP exchanges CE to TG between HDL and other circulating lipoproteins, and thus high CETP activity can lead to low HDL-CE and high HDL-TG (Barter et al., 2003). This can result in a higher TG/CE ratio in HDL, as we found here in the PERI, compared to other menopausal groups. The TG-rich HDL becomes a substrate for hepatic lipase and the concurrent CETP-mediated CE transfer from HDL to other lipoproteins reduces the size of HDL particles (Villard et al., 2012), which agrees with our observation of small perimenopausal HDL. The higher FC/CE- and FC/PL-ratios in the largest XL-HDL particles of the PERI group may suggest lower LCAT activity. LCAT, which esterifies FC to CE, is a key determinant of HDL’s ability to deplete FC from cells (Johnson et al., 1991). Cholesterol efflux capacity is also affected by estrogens carried by HDL, as high estrogen concentration in HDL has been found to enhance the cholesterol efflux capacity (Badeau et al., 2009). On the other hand, we were not able to detect statistically significant differences in cholesterol efflux capacity between menopausal groups. The supraphysiological concentration of estradiol in the experiment of Badeau et al., however, may have a larger effect on the efflux capacity. The menopausal women’s HDL we used, represents normal physiological levels.

Studies on menopausal differences in the HDL lipid species or FA composition are scant rendering our findings novel. We found differences in the relative concentrations of six lipid species and four FAs between the menopausal groups. Of the lipid species with significant group differences, we verified serum FSH concentration to be positively associated with CE 18:3. In addition, estradiol was negatively associated with PC 36:4, PC 38:3, and PC 38:4, concomitantly FSH was positively associated with PC 36:4, LPC 22:5 and negatively with LPC 22:6. We are not aware of previously published studies that have investigated lipid species of isolated HDL particles in association with menopausal status or measured hormones. Previously Tabassum *et al*. (2022) investigated age and sex differences in serum lipid species and included exploratory analyses of menopausal status. They found PC 38:6 to be increased age-related while our finding suggested that PC 38:6 would decrease from premenopause to postmenopause albeit no significant associations were found with hormone levels. The only lipid species Tabassum *et al*. found to be linked to menopause was SM 38:2;2, while we found no significant differences in SM species between menopausal groups. Higher PC 38:6 and LPC 22:6, of which we observed the lowest relative concentration in postmenopausal HDL and negative association with FHS, Cheng *et al*. have linked to better cardiovascular health (Cheng et al., 2015).

Regarding FAs, we mostly observed menopause-status or hormone level-related differences in the low-concentration odd-chain FAs of HDL with decreasing trends from pre- to postmenopause. The odd-chain FAs may be diet-derived or result of gut microbiota metabolism (Jenkins et al., 2017). HDL lipidome has been shown to respond to changes in diet (Zhu et al., 2019), thus it is tempting to assume that the differences we observed could also be attributed to differences in diet. However, our data does not allow us to evaluate detailed differences in the quality and quantity of fats or carbohydrates in the diet, hence this assumption cannot be confirmed. In our data, one even-chain FA, 18:3n-6 (gamma-linolenic acid) was found to be significantly highest in postmenopause. The 18:3n-6 is a precursor of arachidonic acid 20:4n-6, which again is a precursor of eicosanoids including platelet-made blood clotting agent thromboxane, and is linked with systemic inflammation and elevated blood pressure (Sheppe & Edelmann, 2021). We observed a trending increase of the upstream 20:4n-6 from pre- to postmenopause, which also may indicate increased susceptibility to inflammation in postmenopause. The 20:4n-6 also rose up in regression analysis to be negatively associated with estradiol. It is a possible component of PC-species PC 38:4 and PC 36:4, which were negatively associated with estradiol.

ApoA-I is the most abundant protein of HDL and may exist in multiple copies in each particle. The levels of apoA-I and apoA-II are estrogen-responsive; increasing with postmenopausal administration of estrogen therapy and rapidly decreasing after its cessation (Applebaum-Bowden et al., 1989). Therefore, it was not a surprise to observe lower levels of apoA-I and apoA-II in the HDL from peri- and postmenopausal women compared to premenopausal women. ApoA-I is a necessary activator of LCAT to produce spherical HDL while apoA-II-containing HDL particles react less with LCAT (Rye et al., 2003). These apoA-II-rich HDL particles become substrates of hepatic lipase creating small, lipid-poor HDL particles. The apoA-II/apoA-I ratio was the lowest in perimenopause, although only trending. ApoA-II/apoA-I ratio has also been found to indicate the stability of HDL particles as apoA-II prevents apoA-I dissociation from HDL particles (Lund-Katz & Phillips, 2010; Rye et al., 2003). Apart from apoA-I and apoA-II, the other apolipoproteins are exchangeable and present only in a fraction of all HDL particles in circulation. Altogether, we quantified the 11 most abundant apolipoproteins. The size of the HDL particles defines particle curvature, thereby affecting particle cargo as well as the affinity of particles to vascular arterial intima and proteoglycans (Lommi et al., 2011). Some proteins prefer smaller HDL particles, while others prefer larger ones. For example, apoC-III, is found mainly in larger HDL2-particles (Davidson et al., 2009). This agrees with the notion that we found apoA-I-normalized amount apoC-III to be lowest in PERI as the average HDL particle sizes were the lowest in PERI. Also, SAA4, which prefers the small particle size, was found to be the highest in PERI. However, we examined the total HDL pool and could not determine the distinct protein compositions of particles of different sizes.

The high HDL TG proportions and low apoC-I levels we observed in perimenopausal women agree with studies where the low abundance of apoC-I increases plasma TG levels and TG enrichment to HDL (Larsson et al., 2013; Pillois et al., 2012). ApoC-I and apoC-III are inhibitors of lipoprotein lipase (Larsson et al., 2013) and CETP in normolipidemic human subjects (Pillois et al., 2012). ApoC-III and apoE are inversely correlated with estrogen levels (Applebaum-Bowden et al., 1989; Li et al., 2023). Therefore, it is puzzling that the lowest abundances of apoC-III and apoE we found were not in premenopausal women with the highest estrogen concentrations, but in perimenopause where estrogen levels may fluctuate. In addition to the lipid metabolism, ApoE is also involved with immune defense mechanisms (Chernyaeva et al., 2023). In contrast to anti-inflammatory apoE, another HDL apolipoprotein, SAA4 is linked to dysfunctional HDL, leading to low-grade inflammation (de Beer et al., 1995) and susceptibility to deep vein thrombosis (Fernández et al., 2019). SAA isoforms can replace one or more apoA-I from HDL particles and thus influence particle structure and function (Webb, 2021). We observed the highest relative amounts of SAA4 in the PERI group, significant when the protein levels were normalized to apoA-I. This is logical because in our study the perimenopausal state was characterized by the smallest HDL particles and SAA1 and SAA4 are known to preferably exist in the smallest and densest particles (Davidson et al., 2009). This suggests that in perimenopausal women part of the apoA-I may have been replaced with SAA4. The low level of apoE concomitant with increased SAA4 may hence indicate the decreased anti-inflammatory capacity of HDL particles, especially in a perimenopausal state. The higher proportion of pro-inflammatory 20:4n-6 from pre- to postmenopausal groups and the significant negative association with estradiol levels are also in agreement with the decreased anti-inflammatory properties of HDL with progressing menopause.

The single lipid species-level analysis was not efficient in separating menopausal groups. However, the hypothesis of disturbed HDL was supported by the results from our PLS-DA analysis of HDL-multi-omics. A group of HDL-proteins including mainly apolipoproteins and mitochondrial proteins was positively correlated with saturated and mono- and diunsaturated TGs and negatively with polyunsaturated TGs with longer chain FAs. In the heatmap that visualized the data from the menopausal group’s point of view, proteins, and lipids separated the samples of the PERI group from the samples from the PRE and POST groups.

We acknowledge that our study has some limitations. Firstly, this study is an observational study on healthy relatively normolipidemic Caucasian white women without known metabolic diseases or medications. Thus, our results may not be directly generalizable for women representing other ethnic backgrounds or women with different levels of obesity, metabolic syndrome, or otherwise unhealthier conditions. However, to our knowledge, this is the first multi-omics attempt to characterize HDL composition in well-defined menopausal groups, and as such, it has merits albeit being a descriptive rather than a full mechanism-revealing study. Furthermore, we have measured estradiol and FSH concentrations and could therefore inspect if hormone levels are associated with HDL-lipid content. For the association models, we carefully considered potential confounders, such as body adiposity, which significantly contributes to estrogen metabolism before and after menopause (Hetemäki et al., 2017, 2021), and is known to be associated with HDL characteristics. Instead of BMI, we chose to control our models for waist circumference, which is a better estimate of the central body adiposity and has been shown to have equivalent utility for predicting alterations in blood lipids as dual-energy X-ray absorptiometry (Vatanparast et al., 2009). Physical activity, on the other hand, is a known contributor to HDL particle characteristics, with higher activity levels being associated with large HDL particle counts, size, and apoA-1 and HDL-C concentrations (Aadland et al., 2013). Thus, we additionally controlled regression models with physical activity level. Smoking might also influence HDL composition. However, there were only 16 smokers (7.4%) among the 216 women whose samples were used to construct pools for HDL studies. Thus, we considered the potential effect of smoking to be minimal and did not include smoking as a confounder in the analysis. Furthermore, our study did not contain food diaries, hence the detailed diet composition was not available for this study.

Secondly, we did not analyze individual samples, but rather pooled serum samples representing altogether 126 individuals. This was a deliberate choice taken for practical reasons: the HDL isolation protocol is manually laborious and requires a substantial amount of serum for all the designated analyses. Pooling samples decreased the individual variation, which allowed a smaller number of samples per group to be used. This may, alas, also hide some individual findings and mask outliers by causing regression to mean.

Thirdly, HDL particles can be isolated with many different methods each providing compositionally and functionally slightly different HDL particle pools (Holzer et al., 2022). Here, we chose the most common method that has been available since the 50’s, the density gradient ultracentrifugation. We are aware that HDL may shed proteins during ultracentrifugation, and thus the protein content may not be comprehensive. We also detected traces of apoB and apolipoprotein(a) despite of using the two-step sequential ultracentrifugation method, which first separates apoB-containing lipoprotein particles, followed by separation of the ApoA-I-containing particles. ApoB, referring to apoB-100 protein, is an integral part of lipoprotein(a)-particles with a density similar to HDL, thus it is not uncommon to identify these traces in HDL proteomics. Therefore, we consider them as co-eluting contaminants rather than HDL-associated proteins. Due to practical reasons, we were not able to apply the SEC as a second purification step for our HDL isolates. However, as EVs are known to enrich in the same ultracentrifugation fraction with HDL (Simonsen, 2017), we also inspected the EV content of the same pooled sera used for the HDL isolations. Taken that the concentration of EVs has been considered negligible when the HDL properties are described (Nieuwland & Siljander, 2023; Simonsen, 2017) and that no differences were found in CD81 and CD63 positive EVs distribution, although some differences were found in the CD41 and CD9 positive EVs between PRE and POST groups, we consider that the input of EVs on our results is minor.

In summary, we conducted an extensive analysis of HDL particle composition in menopausal women, including size distribution, lipidomics (main lipid classes, lipid species, and fatty acids), and proteomics. Our findings reveal distinct differences in HDL among women in different menopausal stages. Interestingly perimenopause stands out from the premenopause and postmenopause, and notably, perimenopausal women exhibit several cumulative differences in HDL. These differences manifest particularly in particle size, lipid class, and protein compositions: the particles in perimenopause are small and contain a proinflammatory lipid and protein composition. This opens a new focus on the perimenopausal phase as a period of transient changes and warrants further studies, especially to determine whether these observed differences in proinflammatory proteins and lipids are specific to the perimenopausal state with hormonal fluctuations.

## Methods

This study uses serum samples and data from The Estrogenic Regulation of Muscle Apoptosis (ERMA) study (Kovanen et al., 2018). The study was conducted according to the declaration of Helsinki. All participants provided informed written consent, and the study was approved by the Ethical Committee of the Central Finland Health Care District (8U/2014). Among 1393 women who came to the first blood sampling session and had their menopausal status defined based on menstrual diaries and FSH measurements (Kovanen et al., 2018), we selected 216 representative healthy participants (72 women for each menopausal group) with strict exclusion criteria to allow forming 12 serum sample pools representing each pre-, peri- and postmenopausal status. Each pool was constructed from a 1.2 ml serum of six study participants and thereafter aliquoted and stored at -80 °C as three fractions later used for NMR, EV inspections, and HDL purifications.

Exclusion criteria for the current study included cancer, inflammatory diseases (asthma, rheumatoid arthritis, iritis, allergies, celiac disease, psoriasis, rosacea, flu-like symptoms, gastrointestinal disorders), metabolic disorders (diabetes mellitus types 1 and 2, elevated blood pressure, or disorders of lipid metabolism), any cardiovascular diseases, use of hormonal contraceptives or menopausal hormone therapy, statins, and cortisone. In addition, participants with missing information on diseases or BMI were not included.

Leisure-time physical activity and overall quality of the diet were assessed with structured questionnaires as in (Karppinen et al., 2022). The physical activity volume is expressed as metabolic equivalent hours per day (MET-h/d), which accounts for the duration, frequency, and intensity of leisure-time physical activity and the time spent on active commuting. The diet quality was assessed as a sum score computed based on the participant’s responses to a 45-item food-frequency questionnaire. The higher score reflects a healthier diet. BMI was calculated as body mass per height squared (kg/m^2^).

### Blood sampling and serum analysis

Venous blood samples were drawn from the antecubital vein in the supine position, after overnight fasting, between 7 and 10 a.m., in Vacuette® CAT Serum Clot Activator tubes (Greiner Bio-One GmbH, Austria). After a 30-minute incubation at room temperature to allow clotting, the samples were centrifuged at 2200 × g for 10 min at RT. The obtained serum was aliquoted and stored at −80 °C until analysis. Cortisol, insulin, estradiol, sex hormone-binding globulin, and FSH concentrations of serum were determined using IMMULITE 2000 XPi (Siemens Healthcare Diagnostics, UK). Glucose, cholesterol, LDL-C, HDL-C, and TG concentrations were determined using a Konelab 20XTi analyzer (Thermo Fisher Scientific, Finland).

### Nuclear magnetic resonance spectroscopy

The first fraction (0.5 ml) of the pooled serum samples was used to analyze lipoproteins and metabolites with a targeted proton nuclear magnetic resonance (1H-NMR) spectroscopy platform (Nightingale Health Ltd, Finland; biomarker quantification version 2020). The technical details of the method have been reported previously (Soininen et al., 2015).

### Extracellular vesicle characterization

The second fraction (0.6 ml) of pooled serum samples was used to assess EVs. EVs were isolated with SEC. In short, for EV analysis, 500 µl of serum was loaded onto the column (qEV original/70nm, IZON Science, USA), and eluted with dPBS. Four fractions of 500 µl (fractions 1-4) were collected after the void volume (2.7 ml) with an automated fraction collector (IZON Science, USA), combined and concentrated using ultrafiltration (Amicon Ultra, Merck KGaA, Germany). To dilute samples for interferometric *reflectance* imaging sensing (SP-IRIS), particle counts of EVs were measured by nanoparticle tracking analysis using ZetaView (Particle Metrix, Germany) and its corresponding software (version 8.05.12 SP2). Before measurements were taken, the accuracy of the ZetaView was verified using 100 nm polystyrene beads. Samples were diluted in 0.1 μm filtered dPBS to achieve a particle count in the range of 200–500 (1:5000–1:10000). Analyses were conducted in the scattered light mode with a laser wavelength of 488 nm. Next, the isolated EVs were analyzed with single particle SP-IRIS using an ExoView^TM^ Plasma Tetraspanin kit and an ExoView^TM^ R100 scanner (NanoView Biosciences, USA) according to the manufacturer’s instructions. The samples were diluted to about 5 x10^9^ particles /ml with the incubation buffer provided in the kit. All samples (40 µl of each) were added to the chip and incubated at room temperature for 16 h. The samples were then subjected to immuno-fluorescence staining using fluorescently labelled antibodies (CD9/CD63/CD81, provided in the kit), washed, dried, and scanned. Data was analyzed using the NanoViewer analysis software version 3.0.

### Density gradient ultracentrifugation

The third fraction (6 ml) of pooled serum samples was used for HDL particle isolation with density gradient ultracentrifugation. First, the APOB-containing particles were isolated using potassium bromide (KBr) -based density buffer (0.3184 g/ml KBr, PBS, 0.1% EDTA). Volume was set to 9 ml, and the serum was ultracentrifuged for 20 hours in a Ti 70.1 rotor (Beckman Coulter, IN, USA; 65000 rpm; g_max_ = 388024 x g) at a density of 1.063 g/ml. After this, the upper 2.25 ml fraction (i.e., the apoB-containing particles), and middle phase (4.5 ml) were collected from the top of the ultracentrifugation tube and stored at -80 °C. The bottom phase of 2.25 ml was collected, the density was set to 1.21 g/ml with solid KBr, and the volume was reset to 9 ml with isolation buffer. Next, the samples were ultracentrifuged in a Ti 70.1 rotor for 48 hours, after which the top 2.25 ml HDL particle-containing phase, was collected, and stored at -80 °C prior to further analysis.

### Lipid extraction and protein isolation

Stored HDL was thawed on ice, after which the protein component was precipitated with methanol. The lipid content was extracted with sequential extraction with methanol, acetone, diethyl ether ethanol, and chloroform/methanol. Combined supernatants containing the lipid proportion were dried under nitrogen gas (N_2_) flow, reconstituted with chloroform-methanol (1:2) mixture and stored in a glass vial at -20 °C until analysis. The combined protein-containing pellets were dried under N_2_ flow, reconstituted with urea-thiourea buffer (7M urea, 2M thiourea, 0.2M ammonium bicarbonate, 4% CHAPS) and stored at -20 °C until analysis.

### Lipid mass spectrometry

Prior the MS analysis, the sample solutions in chloroform/methanol (1:2 by vol) were spiked with a quantitative standard mixture (SPLASH, Avanti, Alabama, USA), and to enhance ionization and to prevent adduct formation, 1% HN_3_ was added just before the analysis. The mass spectra indicating all positively or negatively charged lipids, lipid precursors of the ions m/z 184 (LPCs, PCs and SMs), m/z 369 (CEs) and m/z 241 (phosphatidylinositols), and lipids assigned through neutral loss of amu 141 (phosphatidylethanolamines) and 87 (phosphatidylserines) were recorded on an Agilent 6410 Triple Quad LC-MS after infusing the sample solution with a syringe pump (flow rate 10 µl/min). The data were further analyzed with MassHunter Qualitative Analysis Navigator B.08.00 (Agilent Technologies, CA, USA) and LIMSA software (Haimi et al., 2006).

### Fatty acid gas chromatography

After removing the organic solvents with N_2_ flow, aliquots of the extracted lipids were converted to FAMEs by heating in 1% methanolic H_2_SO_4_ with hexane as cosolvent at 95 °C for 120 min under N_2_ atmosphere. Following water addition, the FAMEs were extracted with hexane, dried with anhydrous Na_2_SO_4_, concentrated under N_2_ flow, and stored at -80 °C. The FA composition of the HDL particles was analyzed by injecting the FAME samples into a GC-2010 Plus gas chromatograph (Shimadzu Scientific Instruments, Japan) with a flame-ionization detector and integrating the resulting chromatographic peaks using a Shimadzu GC solution 2.42 software. Identification of the FAMEs was based on EI mass spectra recorded by a GCMS-QP2010 Ultra (Shimadzu Scientific Instruments) with a mass selective detector. Both systems were equipped with Zebron ZB-wax capillary columns (30 m, 0.25 mm ID and film thickness 0.25 µm; Phenomenex, CA, USA). The FA compositions were expressed as mol-% profiles.

### Label-free HDL proteomics and bioinformatic data analysis

The protein samples were digested in Amicon Ultra 0.5 centrifugal filters using a modified FASP method, as described (Scifo et al., 2015). The peptides were cleaned using C18-reverse-phase ZipTip^TM^ (Millipore, Merck KGaA). The dried peptide digest was re-suspended in 1% trifluoroacetic acid and sonicated in a water bath for 1 min before injection. Protein digests were analyzed using nano-LC-Thermo Q Exactive Plus. The peptides were separated by an Easy-nLC system (Thermo Scientific) equipped with a reverse-phase trapping column Acclaim PepMap^TM^ 100 (C18, 75 μm × 20 mm, 3 μm particles, 100 Å; Thermo Scientific). The injected sample analytes were trapped at a flow rate of 2 µl min^-1^ in 100% of solution A (0.1 % formic acid). After trapping, the peptides were separated with a linear gradient of 120 min comprising 96 min from 3% to 30% of solution B (0.1% formic acid/80% acetonitrile), 7 min from 30% to 40% of solution B, and 4 min from 40% to 95% of solution B. Each sample run was followed by an empty run, which comprises a trap column flush with 16 µl of 80% acetonitrile to reduce the sample carryover from previous runs. LC-MS data acquisition was done with the mass spectrometer settings as follows, the resolution was set to 140,000 for MS scans, and 17,500 for the MS/MS scans. Full MS was acquired from 350 to 1400 m/z, and the 10 most abundant precursor ions were selected for fragmentation with 30 s dynamic exclusion time. Ions with 2+, 3+, and 4+ charges were selected for MS/MS analysis. Secondary ions were isolated with a window of 1.2 m/z. The MS AGC target was set to 3 x 10^6^ counts, whereas the MS/MS AGC target was set to 1 x 10^5^. Dynamic exclusion was set with a duration of 20 s. The NCE collision energy step was set to 28 kJ mol^−1^.

Following LC-MS/MS acquisition, for the relative quantification of proteins, the raw files were qualitatively analyzed by Proteome Discoverer (PD), version 2.4 (Thermo Scientific). The identification of proteins by PD was performed against the UniProt Human protein database (UniProt release 2020_12 downloaded on January 26^th^, 2021, with 20406 entries) using the built-in SEQUEST HT engine. The following parameters were used: 10 ppm and 0.02 Da were mass tolerance values set for MS and MS/MS, respectively. Trypsin was used as the digesting enzyme, and up to two missed cleavages were allowed. The carbamidomethylation of cysteines was set as a fixed modification, while the oxidation of methionine and deamidation of asparagine and glutamine were set as variable modifications. The false discovery rate was set to less than 0.01 and a peptide minimum length of six amino acids. Chromatograms were automatically aligned by the Progenesis QI for Proteomics^TM^ (NonLinear Dynamics) software using the default values by following the wizard, and those that had an alignment score >67% to the reference run were selected for further analysis. As a final filtering step, only proteins that were detected in at least two technical replicates of two samples in each menopausal group were accepted for further analysis.

### LCAT activity assay

LCAT activity was measured from the serum samples of 10 premenopausal, 9 perimenopausal, and 9 postmenopausal women who were BMI-matched to the mean of the women used in earlier serum pools, i.e., to BMI about 25 kg/m^2^. LCAT activity assay kit (Sigma-Aldrich MAK107, Merck) was used according to the manufacturer’s instructions.

### THP-1 macrophages and cholesterol efflux assay

The THP-1 monocyte line (TIB-202, ATCC, VI, USA) was maintained according to the provider’s instructions. Briefly, monocytes were cultured in RPMI-1640 (Gibco, UK) with 10% heat-inactivated fetal bovine serum (FBS, Gibco) and 0.05 mM 2-mercaptoethanol (Sigma-Aldrich, Merck). For differentiating to macrophages, monocytes were seeded with 100 nM of phorbol 12-myristate 13-acetate (Sigma-Aldrich, Merck) into a 96-well tissue culture plate (60,000 cells/well, 100 µl/well) and incubated for 72 h. The cholesterol efflux assay was performed according to the kit instructions (Sigma-Aldrich MAK192, Merck) without FBS present. Macrophages were incubated with fluorescently labelled cholesterol for 1 h and then equilibrated overnight. After washing with FBS-free RPMI-1640, three concentrations of PRE-, PERI-, and POST-HDL samples (50, 125 and 200 µg/ml total HDL protein) with positive and negative controls were added to the cells and incubated for 4 h. Then, the amount of fluorescence in the supernatant and solubilized cells was measured with a Glomax multi + detection system (Promega, WI, USA). The efflux capacity was calculated from the ratio of fluorescent intensity in the supernatant and the total fluorescent intensity in the supernatant and cell lysate. Cells were kept in a humidified CO_2_ atmosphere at 37°C during culturing, differentiation and efflux assay and passage number 6 was used in efflux assay.

### Statistical Analyses, Data Processing, and Figures

Calculation of statistical significance was performed with IBM SPSS statistics (Version 26, IBM, USA). One-way ANOVA was used with Tukey’s Honest Significant Difference test as a *post-hoc* to explore differences between multiple groups’ means. For skewed variables indicated by histograms and Q-Q-plots, the Kruskal-Wallis test was used. For the LCAT activity test, the paired samples T-test was used to analyze differences between groups. P values smaller or equal to 0.05 were considered significant. Because large correlations were assumed between lipid or protein species, multiple testing corrections were expected to give misleading results and were not done (García-Pérez, 2023; Rothman, 2014).

To investigate whether the three menopausal groups differ by some combination of multiple features, dimensionality reduction with Partial Least Squares Discriminant Analysis (PLS-DA) was performed with R package *mixOmics* (Rohart et al., 2017). As the protein and lipid data were already normalized, no further pre-processing steps were done. To detect correlations between proteins and lipids, omics integration was done with mixOmics. Correlation circles obtained from block sparse PLS-DA were plotted and corresponding Clustered Image Maps (CIM) were calculated. The CIM was re-plotted with a heatmap.2 function of R package gplots to control dendrogram ordering.

Before calculating standardized regression coefficients, dependent and independent variables were first converted into z-values by subtracting the mean and dividing by standard deviation. Multiple linear regression was done in R. Eight models were calculated:

Model 1: independent variable ∼ Estradiol

Model 2: independent variable ∼ Estradiol + waist circumference

Model 3: independent variable ∼ Estradiol + leisure time physical activity

Model 4: independent variable ∼ Estradiol + waist circumference + leisure time physical activity

Model 5: independent variable ∼ FSH

Model 6: independent variable ∼ FSH + waist circumference

Model 7: independent variable ∼ FSH + leisure time physical activity

Model 8: independent variable ∼ FSH+ waist circumference + leisure time physical activity

The resulting standardized regression coefficients are given in supplementary Tables S3a and S3b. The original data for TG 54:03 had 7 values missing, so the results were calculated without the missing values. Other variables did not have missing data.

Figure 1 was created with BioRender.com. Other figures (except Figure 3a) were plotted with the R package *ggplot2*. The data for the figures was processed using RStudio and R version 4.3.1.

## Data availability

The data that supports the findings of this study are available in the supplementary material of this article. However, the individual-level data is not completely publicly available due to privacy or ethical restrictions. The metadata of the ERMA study is publicly available (doi:10.17011/jyx/dataset/83491). The main code used to conduct this study is available on GitHub at <u>LaakkonenLab/MATCH</u> (github.com).

## Acknowledgments

This study was funded by grants from the Academy of Finland (grant numbers 33028, 309504, 314181, and 335249 to E.K.L). We acknowledge the following core units of the University of Helsinki: the EV Core for SEC and SP-IRIS analyses, HiLIPID of HiLIFE and Biocenter of Finland for lipidomic analyses and the Meilahti Clinical Proteomics unit of the HiLIFE supported by Biocenter Finland for proteomic analyses. We thank the staff at the Faculty of Sport and Health Sciences of the University of Jyväskylä for their invaluable assistance with data collection. We also thank the women who participated in the ERMA study for donating their time and effort.

## Author contributions

E.K.L. is the PI of the ERMA study and together with M.L. designed and supervised the current study. E.K.L. provided funding for the study. S.L. did most of the laboratory work including HDL particle isolations, lipid and protein extractions, ESI-MS, and LCAT assays. S.L. analyzed lipidomics datasets concerning lipid classes and lipid species. H.R. and R.K. performed GC-FID and GC-MSD analyses of fatty acids. R.S. did protein LC-MS and analysis of the data which M.La. supervised. S.L. did data curation for lipidomic and proteomic datasets. E.L. did cholesterol efflux assays. M.P. and P.S. were responsible for the EV isolations and analysis. S.L. constructed the data files and did the statistics and visualization based on them. T-M.K. performed the bioinformatic analyses for Figure 5 and the regression analyses and polished the figures using R packages. S.L. wrote the original draft of the manuscript. T-M.K., M.L., and E.K.L. contributed to the editing of the manuscript. All co-authors have participated in the interpretation of the results and critically commented on the manuscript during the writing process. SL and E.K.L. prepared the final version of the manuscript, which all the co-authors approved.

## Competing interests

The authors declare no competing interests.

## Notes

### Competing Interest Statement

The authors have declared no competing interest.

### Summary of Updates

This version of the manuscrip has been revised by adding some new analyses and updating figures and supplemental tables were accordingly.

