## Supplementary figures for "The lipidome and proteome of high-density lipoprotein are altered in menopause"

**Supplementary Figure S1**

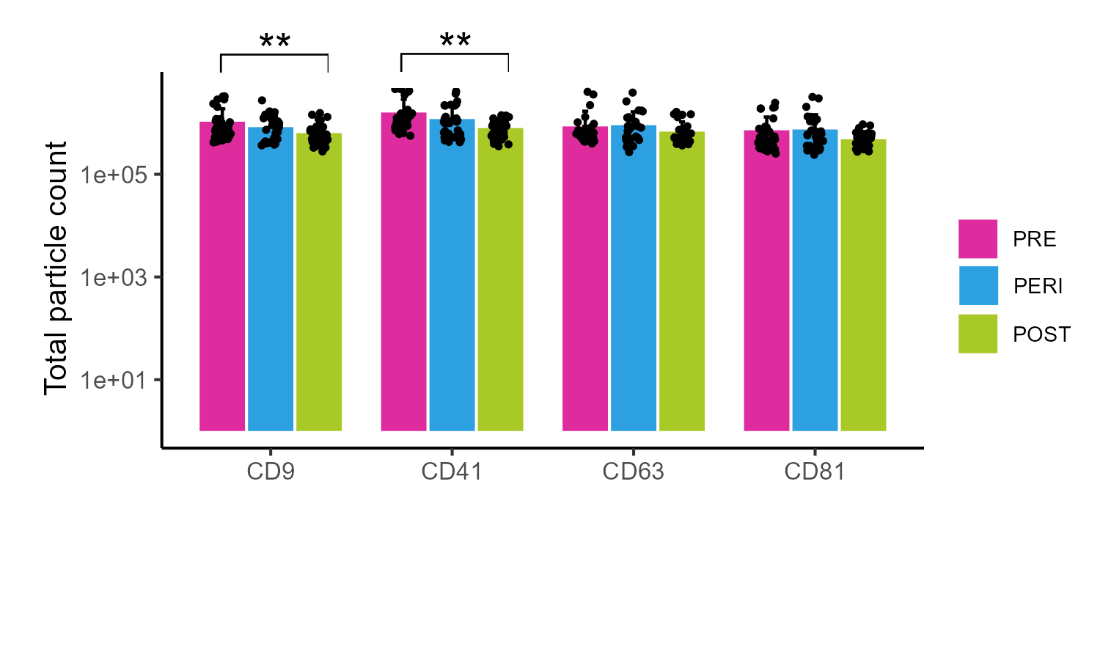

**Supplementary Figure S1. Characterization of the size exclusion chromatography -purified extracellular vesicles (EV).** Distribution of CD9, CD41, CD63 and CD81 positive EVs across menopausal groups. Statistical significance **<0.01. Abbreviations: PRE, premenopause; PERI, perimenopause; POST postmenopause.

**Supplementary Figure S2**

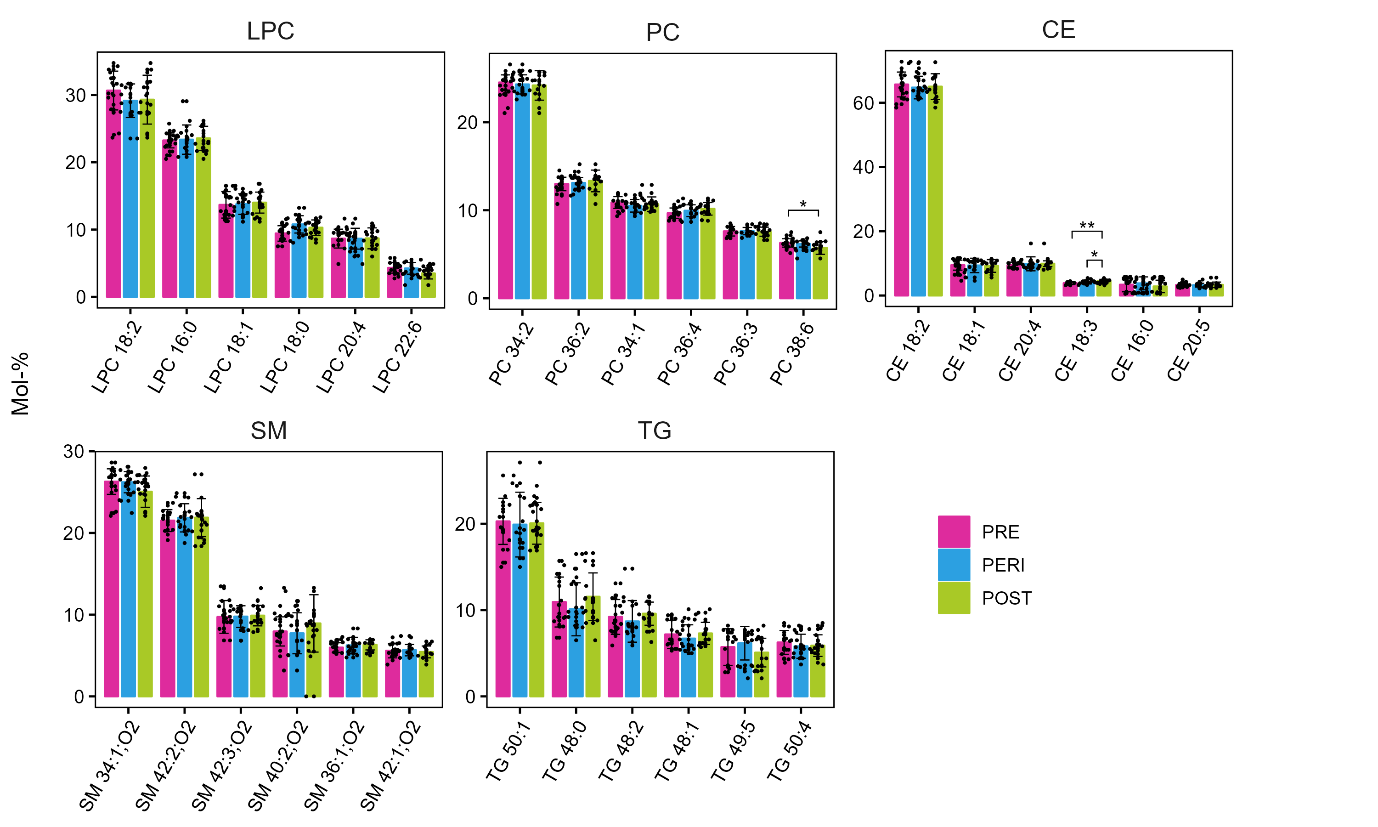

**Supplementary Figure S2.** **The most abundant lipid species observed in the five main lipid classes of the HDL particles.** Lipid species abundances are expressed as mol-% within each lipid class. Lipids were extracted from ultracentrifugally isolated HDL and analyzed with ESI-MS. Statistical significance * < 0.05, **<0.01. Abbreviations: HDL high-density lipoprotein, PRE premenopause, PERI perimenopause, POST postmenopause, LPC lysophosphatidylcholine, PC phosphatidylcholine, CE cholesteryl ester, SM sphingomyelin, TG triacylglycerol.

**Supplementary Figure S3**

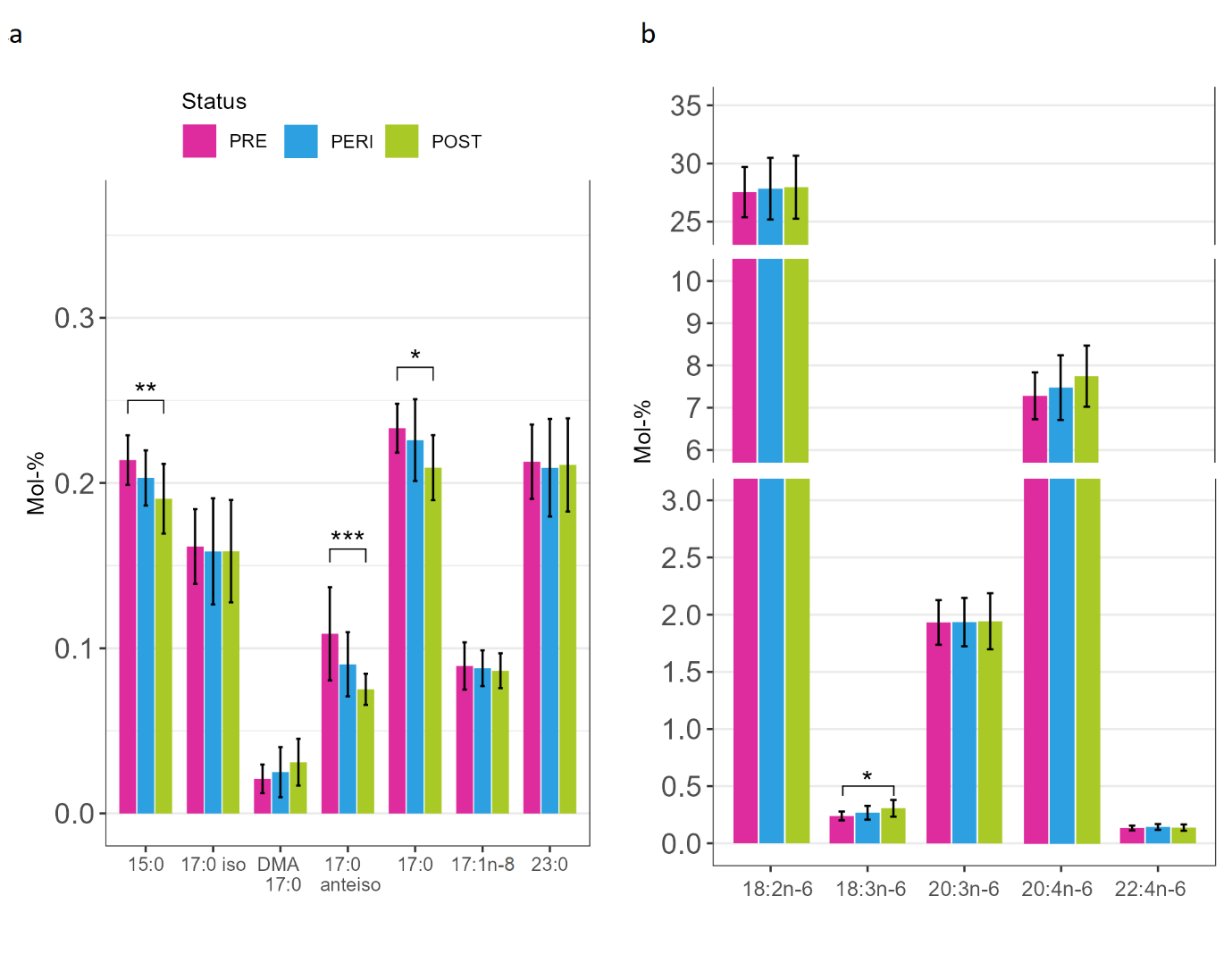

**Supplementary Figure S3.** **Fatty acid composition of HDL lipids in three different menopausal statuses.** (**a)** Odd-chain fatty acids of HDL. **(b)** n-6 pathway fatty acids of HDL. Fatty acids were analyzed with GC-FID and GC-MS. Statistical significance * < 0.05, **<0. 01, ***<0.001. Abbreviations: HDL, high-density lipoprotein; PRE, premenopause; PERI, perimenopause; POST, postmenopause; DMA, dimethyl acetal (represents an alkenyl chain from plasmalogen phospholipids, and is formed when methylating fatty acids for gas chromatography); GC-FID, gas chromatography with flame ionization detector; GC-MS gas, chromatography-mass spectrometry.

**Supplementary Figure S4**

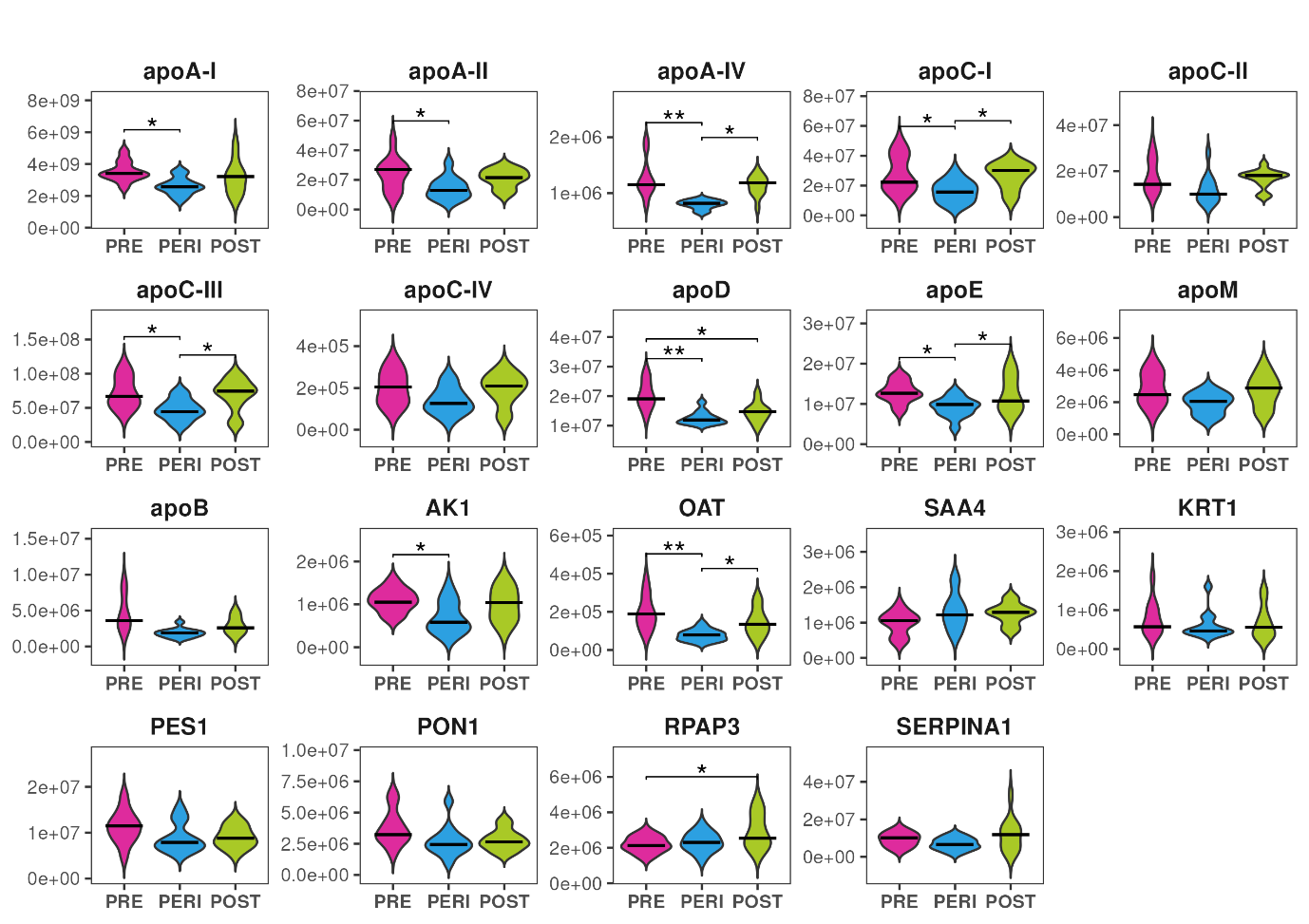

**Supplementary Figure S4.** **Violin plots of unnormalized HDL protein abundances (raw data).** All protein abundances were measured with LC-MS method from the isolated HDL particles. Statistical significance * < 0.05, **<0.01. Abbreviations: PRE premenopause, PERI perimenopause, POST postmenopause, HDL high-density lipoprotein, and LC-MS liquid chromatography-mass spectrometry.

**Supplementary Figure S5**

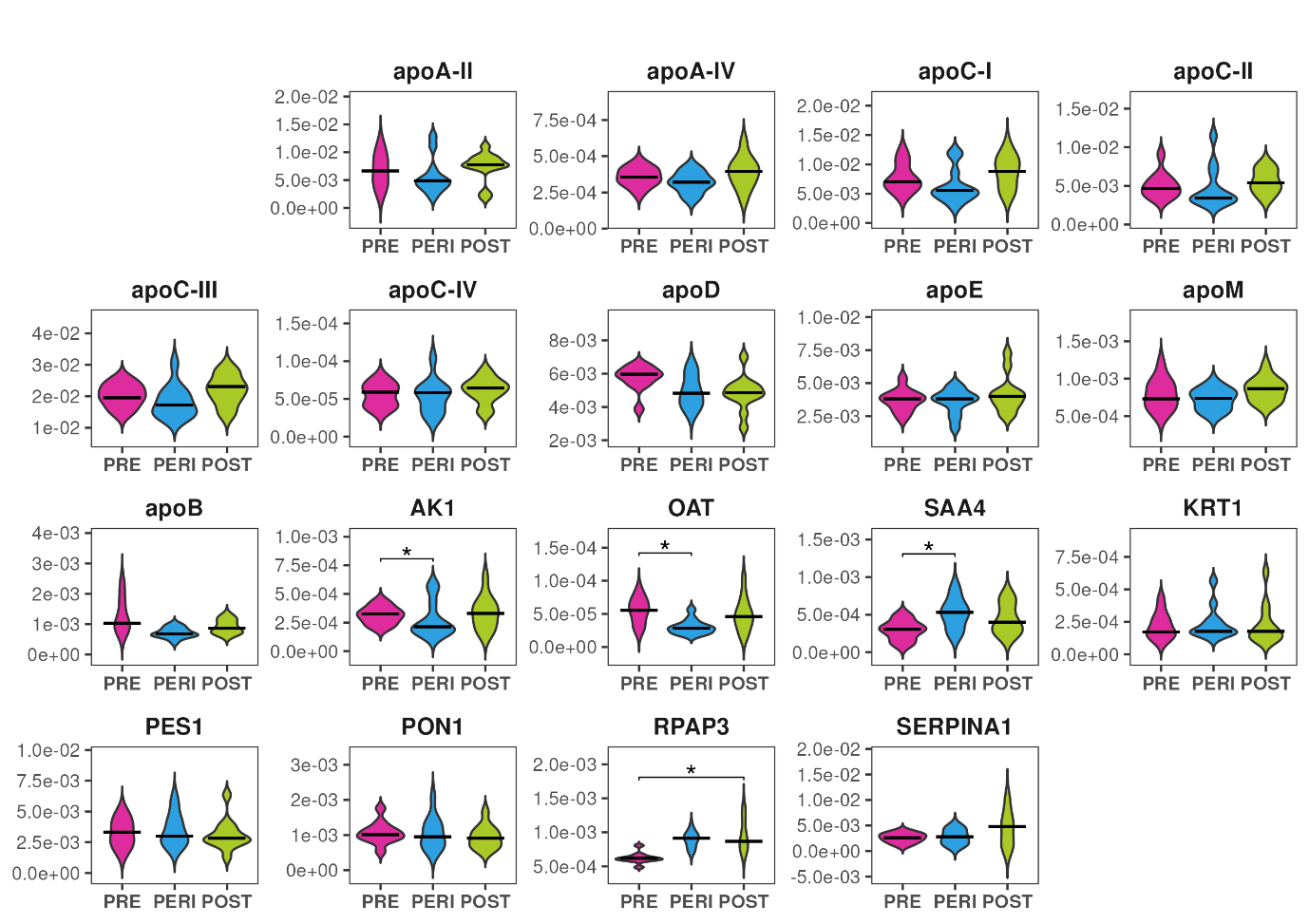

**Supplementary Figure S5. Violin plots of HDL protein abundances normalized to apoA-I.** All protein abundances were measured with LC-MS method from the isolated HDL particles. Statistical significance * < 0.05, **<0.01. Abbreviations: PRE premenopause, PERI perimenopause, POST postmenopause, HDL high-density lipoprotein, and LC-MS liquid chromatography-mass spectrometry.

**Supplementary Figure S6**

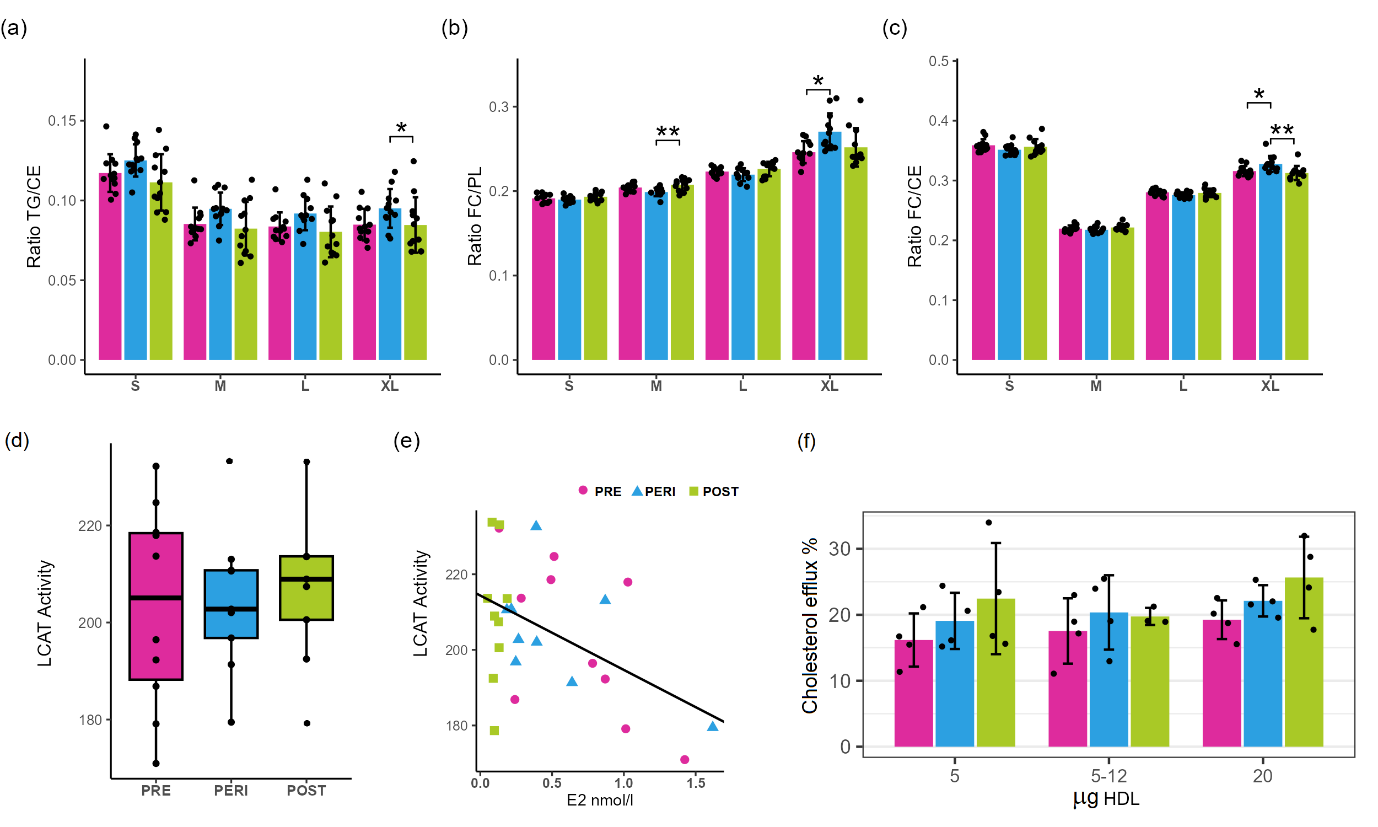

**Supplementary Figure S6.** Functional assessments of the HDL particles across menopausal stages. (a). The ratio of lipid classes TG/CE in HDL particle size categories. (b). The ratio of lipid classes FC/PL in HDL particle size categories. (c) The ratio of lipid classes FC/CE in HDL particle size categories. All lipid class ratios were calculated from lipid classes measured with NMR. (d) LCAT activity assay with serum samples representing PRE (n = 10), PERI (n = 9) and POST (n = 9) women. (e) Correlation plot of the measured LCAT activity and estradiol concentration. (f) Cholesterol efflux assay with 5 µg, 12.5 µg and 20 µg of purified HDL originating from the pooled serum samples of PRE, PERI and POST women. Statistical significance * < 0.05, **<0.01. Abbreviations: HDL high-density lipoprotein, PRE premenopause, PERI perimenopause, POST postmenopause, TG triglycerides, CE cholesteryl ester, FC unesterified cholesterol, PL phospholipid, NMR nuclear magnetic resonance, LCAT lecithin:cholesterol acyltransferase, and E2 17β-estradiol.
